# miniODP: a reusable framework for building multi-omics resources in understudied organisms

**DOI:** 10.1101/2024.01.06.573815

**Authors:** Hang Yang, Zehui Wang, Zhaojun Shan, Hanqiao Shang, Puxuan Jiang, Yingshu Li, Qiang Tu

## Abstract

Building multi-omics resources for understudied organisms requires assay selection, public-data curation, gene identifier handling, portal deployment, and visualization, yet these tasks are rarely packaged into a reusable framework. To fill this gap, we developed the mini Omics Data Portal (miniODP), which combines species pages, gene-centric modules, genome browsing, and sequence search with configuration files, species onboarding workflows, and demonstration data for self-deployment. Alongside the software, we propose miniENCODE core assays: a reduced RNA-seq, ATAC-seq, and H3K27ac profiling set for regulatory analysis. Current miniODP includes seven species, covering 3,568 bulk runs, 3.61 million cells, and 1,865 genome-browser tracks. Matched core-assay datasets support regulatory outputs. As a zebrafish benchmark, using datasets from three core assays and 17 samples, we identified 52,350 enhancer-like signatures (ELSs) and constructed ELS-to-gene linkages and gene regulatory networks (GRNs). Genes near H3K27ac-supported ELSs were expressed at higher levels than those near ATAC-only distal peaks. Literature curation of top TFs from zebrafish and cattle GRNs found 19 direct, 15 indirect, and six unsupported cases among 40 candidates. Datasets lacking matched core assays still provide browsing, visualization, and search functions. miniODP serves as a reusable framework for constructing and extending multi-omics resources for understudied organisms.

**Graphical abstract:** 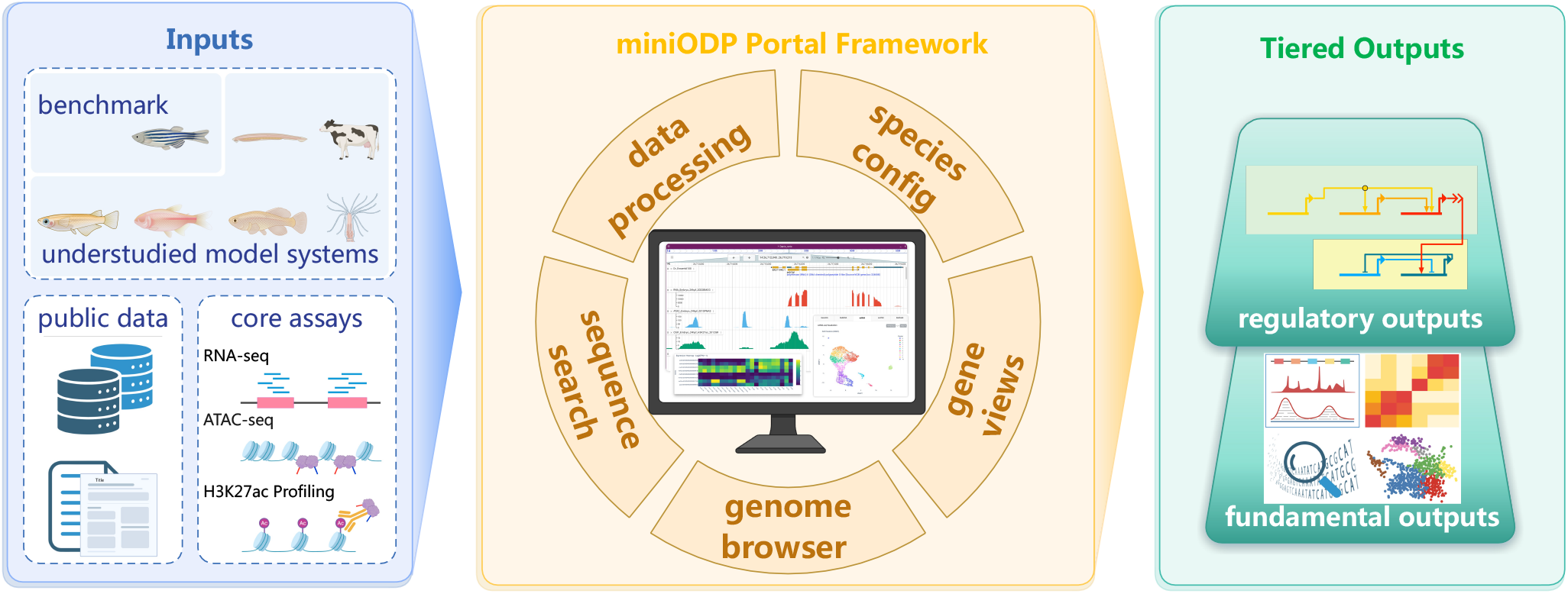

## Introduction

Large-scale functional genomics projects have greatly improved regulatory annotation from genome sequences. ENCODE mapped diverse functional genomic elements in humans and mice, and modENCODE expanded related efforts to *C. elegans* and *D. melanogaster* (1–5). However, most communities working on other organisms cannot replicate this scale. In GEO, about 80% of samples are from humans and mice, the next 9% of samples come from eight species, and the remaining 11% come from more than 6,000 species (Supplementary Figure 1). For many emerging or understudied organisms, the bottlenecks go beyond data generation and include assay selection, public-data curation, gene identifier mapping, portal deployment, and long-term maintenance of these resources.

Some communities have already started tackling these issues. The zebrafish community is a good example and has established multiple platforms. ZFIN is a curated knowledge base that has long served as the community’s central hub (6). Yang et al. and DANIO-CODE provide extensive epigenomic resources (7– 9). Multiple single-cell or multimodal profiling resources have been established, including DanioCell, ZSCAPE, ZebraHub, and developmental chromatin-accessibility maps (10–13). In the meantime, multi-species portals are emerging, including iFish for fishes, DOO for deep-ocean organisms, and MedakaBase for *Oryzias* species (14–16). These platforms offer browsing, search, and visualization functions. However, they generally do not provide the full workflow for building and deploying such resources for other species from scratch.

Therefore, a reusable framework is required for building omics portals in more species. Such a framework should cover both data processing and portal operation. On the data side, heterogeneous inputs have to be converted to expression matrices, genome browser tracks, sample registration, sequence search databases, and gene identifier mapping tables. On the portal side, species pages, gene-centric views, genome-centric views, sequence searching, links between modules, config templates, and deployment documents are necessary. The framework should also be easy to adapt to new species. In addition, it must handle incomplete datasets: a new species could begin with just a genome sequence and some bulk expression data, while advanced species might already have matched RNA-seq, ATAC-seq, and epigenomic data that support ELS and GRN analysis.

To fill the gap, we developed miniODP as a deployable, reusable multi-omics framework. It combines Hugo species information pages, a Python Dash gene-centric module, a JBrowse 2 genome browser, and SequenceServer sequence search. Species-specific behavior is handled by configuration files and species adapters, so the portal can support new species with limited adjustment. We propose a small set of core assays, RNA-seq, ATAC-seq, and H3K27ac profiling, as a resource-efficient starting point for regulatory analyses. We term this assay set miniENCODE. More assays like long-read RNA-seq, Micro-C, DNA or histone modifications, and single-cell omics can be added gradually when needed.

We set up a miniODP instance containing seven species. Using a dataset of three matched core assays and 17 developmental or adult tissue samples from zebrafish, we generated 52,350 enhancer-like signatures (ELSs), ELS-to-gene linkages, and GRNs. We tested selected ELSs using F0 reporter assays. We showed that genes near H3K27ac-supported ELSs are expressed at higher levels than genes near ATAC-only distal peaks, indicating that adding H3K27ac provides a modest but stable information gain. Literature curation of top TFs from zebrafish and cattle GRNs provides direct, indirect, and unsupported evidence. Without matched core-assay data, the portal can still support browsing, search, and visualization, as shown by medaka and killifish. Finally, we compared miniODP with relevant resources to show that it is a reusable framework rather than a fixed species-specific database.

## Materials and Methods

### Zebrafish husbandry

Zebrafish of the TU strain used in this study were raised and bred following standard fish-keeping procedures (17). The fish were maintained in a recirculating aquaculture system with freshwater at 28 °C and a 14-hour light/10-hour dark photoperiod.

### Sample collection

For ovary isolation from adult female zebrafish, we first anesthetized the fish with Tricaine methanesulfonate (MS222) and positioned them in a 2% agarose groove. Using ophthalmic scissors and fine forceps, we performed abdominal dissection and transferred the ovaries to a tube containing 2 mL of DPBS (Gibco, cat. C14190500BT). The ovarian tissue was mechanically dissociated into <1 mm fragments using microdissection scissors. After tissue sedimentation, we replaced the medium with 2 mL of digestive enzyme mixture I (3 mg/mL collagenase I (Worthington, cat. LS004196), 3 mg/mL collagenase II (Worthington, cat. LS004174), and 1.6 mg/mL hyaluronidase (Worthington, cat. LS002594) in Leibovitz L-15 Medium (Gibco, cat. 21083027)). The mixture was incubated at 37 °C with gentle agitation at 15 rpm on an orbital rotator. We monitored dissociation every 10 minutes until the tissue was fully dissociated. After centrifugation at 300 *g* for 5 minutes, the cell pellet was resuspended in 2 mL of digestive enzyme mixture II (0.25% TRYPSIN-EDTA (Gibco, cat. 15400-054), 0.8 U/mL DNase I (Invitrogen, cat. M0303S) in L-15) and incubated for 10 minutes at 37 °C. The enzymatic reaction was stopped by adding 200 µL of FBS (Sigma-Aldrich, cat. F8318). The cells were then washed twice with 1 mL of DPBS containing 0.04% BSA, centrifuged at 500 *g* for 5 minutes, and filtered through a 100 µm cell strainer.

For embryo collection, we followed established staging criteria (18) and verified developmental stage through morphological examination. Chorions were removed by incubating embryos in 1 mg/mL Pronase for 10 minutes followed by DPBS washing. Yolks were removed using 1 mL of deyolking buffer (55 mM NaCl, 1.8 mM KCl, and 1.25 mM NaHCO_3_) with 20-30 pipetting cycles. Embryonic tissues were homogenized in 2 mL of FACSmax cell dissociation solution (Genlantis, cat. T200100) for 5 minutes at room temperature. The cell suspension was filtered through a 40 µm cell strainer and centrifuged at 310 *g* for 5 minutes. Cell density and viability were quantified using the Countstar Rigel S2 system.

### Library preparation

#### ATAC-seq library

Freshly dissected samples were immediately processed for ATAC-seq. Tissues were resuspended in 1.5 mL of lysis buffer (10 mM Tris-HCl [pH 7.4], 10 mM NaCl, 3 mM MgCl_2_, 0.10% Tween-20 [Bio-rad, cat. 1662404EDU], 0.10% NP-40 [Abcam, cat. ab142227], 0.01% Digitonin, 1% BSA in nuclease-free water). After filtration through a 40 µm cell strainer, nuclei were collected by centrifugation at 500 g for 5 minutes. ATAC-seq libraries were prepared following established protocols (19). Nuclei were resuspended in 50 µL of transposition mix (10 µL 5x TTBL, 5 µL TTEmix V50 [Vazyme, cat. TD501], 0.5 µL 1% digitonin, 0.5 µL 10% Tween-20, 34 µL ddH_2_O). After PCR amplification, libraries were size-selected using 0.6x AMPure beads (Beckman, cat. A63880) to remove large fragments, followed by 1.2x AMPure beads for library recovery. All ATAC-seq libraries were sequenced on the Illumina NovaSeq platform, with two biological replicates per sample and a minimum of 60 million paired-end 150 bp reads.

#### CUT&Tag library

CUT&Tag libraries were prepared following established protocols (20) using the Hyperactive Universal CUT&Tag Assay Kit for Illumina (Vazyme, cat. TD903). Cells were harvested and washed twice with 1.5 mL of Wash Buffer using gentle pipetting. 10 µL of activated concanavalin A-coated magnetic beads were added and incubated at room temperature for 10 minutes. The bead-bound cells were resuspended in 50 µL of chilled Antibody Buffer containing either 1 µL of H3K27ac antibody (Abcam, cat. ab4729) or Rabbit IgG (Abcam, cat. ab171870) as primary antibody, followed by overnight incubation at 4 °C.

After removing the primary antibody using a magnet stand, cells were incubated with Guinea Pig anti-Rabbit IgG (Heavy & Light Chain) antibody (Antibodies-Online, cat. ABIN101961) at room temperature for 60 minutes. The cells were then washed twice with 0.2 mL of Dig-Wash buffer for 5 minutes each using the magnet stand. For tagmentation, 50 µL of 0.04 µM pA/G-Tn5 adapter complex in Dig-300 Buffer was added with gentle vortexing and incubated at room temperature for 1 hour. After additional washes with Dig-300 Buffer, cells were resuspended in 50 µL of Tagmentation buffer (10 µL of 5x TTBL in 40 µL of Dig-300 Buffer) and incubated at 37 °C for 60 minutes.

DNA was extracted by adding 5 µL of Proteinase K, 100 µL of Buffer L/B, and 20 µL of DNA Extract Beads to 50 µL of sample. After two washes with 200 µL of Buffer WA and WB, the beads were dried and resuspended in 22 µL of nuclease-free water. Libraries were amplified by mixing 15 µL of DNA with 5 µL of universal i5 primer, 5 µL of uniquely barcoded i7 primer, and 25 µL of 2x CAM. Final libraries were purified using 100 µL of VAHTS DNA Clean Beads (Vazyme, cat. N411).

### Enhancer reporter assays

To test selected heart-associated ELSs, we performed *Tol2* transposon-mediated F0 zebrafish reporter assays following established protocols (21). Candidate ELS sequences were PCR amplified from zebrafish genomic DNA using primers listed in Supplementary Table 2 and cloned into the pSTG vector upstream of a minimal EF1a promoter-EGFP cassette. The resulting reporter plasmids were co-injected with transposase mRNA into one-cell-stage TU zebrafish embryos. Embryos were examined at 3 dpf using fluorescence microscopy (Leica), and at least 100 embryos were scored for each construct. Constructs with GFP signal in at least 5% of scored embryos were treated as having detectable reporter activity. Because these were F0 transient assays, reporter activity was interpreted as support for selected candidate ELSs rather than as stable-line validation of the full ELS set. The 23 tested constructs were prioritized from the heart-specific ELS subset by ATAC-seq and H3K27ac signal strength.

### Data analysis

#### Data curation and processing

We constructed miniODP as a seven-species multi-omics resource covering zebrafish, medaka, turquoise killifish, Mexican tetra, Hydra, lancelet, and cattle. Detailed data sources and references are provided in Supplementary Table 3. The portal-level resource includes species with different data completeness. Matched RNA-seq, ATAC-seq, and H3K27ac data were used for ELS-centered analyses when available, and selected samples with suitable expression, chromatin, and motif inputs were used for GRN construction; species lacking sufficient H3K27ac coverage were integrated into the portal through the available expression, single-cell, genome-browser, or sequence-search modules.

In portal statistics, bulk runs refer to bulk omics sequencing runs included in miniODP, usually corresponding to run-level accessions such as SRR, ERR, or DRR records. Single-cell sample groups refer to curated public matrix-level display units in Dash; each group can represent one or more source libraries or runs merged during curation.

Raw sequencing data were inspected with FastQC (https://www.bioinformatics.babraham.ac.uk/projects/fastqc/) and filtered with Trim Galore (http://www.bioinformatics.babraham.ac.uk/projects/trim_galore/) when reprocessing from raw reads was required. Processed public matrices or tracks were retained when they were the primary release format of a source study and were compatible with the miniODP data model.

For RNA-seq datasets reprocessed from raw reads, reads were aligned to the species-specific reference genomes used by the corresponding miniODP species modules using HISAT2 (version 2.2.1) (22), followed by gene quantification with Stringtie (version 2.1.2) (23). Gene expression levels were normalized and reported as transcripts per kilobase million (TPM).

For ChIP-seq, CUT&Tag, and ATAC-seq analysis, reads were aligned to the corresponding species reference genomes using Bowtie2 (version 2.4.2) (24). We removed duplicate fragments using SAMtools (25) and Picard (http://broadinstitute.github.io/picard/), called peaks with MACS2 (version 2.2.7.1) (26), and indexed resulting alignments using SAMtools. Epigenomic quality control included insert-size distribution, TSS enrichment score, chromosome-level mapping summaries, and Pearson correlation between replicates.

For methylation analysis (WGBS and Bisulfite-Seq), we processed data using Bismark (version 0.22.3) (27) and deepTools (version 3.4.3) (28). Coverage files in BigWig format are available through JBrowse 2.

Because miniODP integrates multiple assay types and public studies released in different formats, we did not require all datasets to pass through a single assay-specific workflow such as nf-core. Instead, assay-specific scripts and a containerized toolchain produced shared portal-ready outputs, including expression matrices, peak and coverage files, ELS calls, GRN files, single-cell objects, metadata tables, and genome-browser or BLAST assets.

We analyzed single-cell RNA-seq data using Seurat (version 3.2.2) (29), with input from either pre-processed gene expression count tables or raw data processed through Cell Ranger (version 4.0.0). Data sources are listed in Supplementary Table 3. We applied standard quality filters based on the number of detected features, total counts per cell, and mitochondrial content. For snATAC-seq data, we used gene activity score matrices to estimate gene expression levels from chromatin accessibility patterns.

We reduced dimensionality with principal component analysis (PCA) on the top 2000 most variable genes and selected the number of principal components using ElbowPlot and JackStrawPlot analyses. Cell clusters were visualized using UMAP and t-SNE. We analyzed differential gene expression between clusters using the FindAllMarkers function with default parameters.

#### miniODP platform implementation

The public miniODP portal combines Hugo species pages (https://gohugo.io/), gene-centric applications built with the Python Dash framework (https://dash.plotly.com/), JBrowse 2 genome browser sessions (30), and SequenceServer-based sequence search (31). Hugo stores species-level display information, default module links, and summary statistics. The Dash applications provide gene information, bulk RNA, scRNA, scATAC, and BulkMulti modules when the corresponding data are available. JBrowse 2 stores genome assemblies and multi-omics tracks. SequenceServer provides BLAST searches against genome, cDNA, and protein databases.

Species-specific behavior is configured rather than hard-coded in shared application logic. Species display settings are managed through TOML configuration files, while Dash species adapters define gene identifier rules, search fields, and annotation links. The current adapter system supports Ensembl-style identifiers, dual identifier systems, and local gene ID schemes. Depending on available annotations, gene search can use gene identifiers, gene names, GO terms, InterPro domains, TF families, or orthologs; gene pages can link to Ensembl (32), AnimalTFDB (33), and InterPro (34) where appropriate.

Data directories are organized by species and module, with conversion scripts preparing gene information, bulk expression tables, single-cell objects, genome browser tracks, and BLAST database manifests. Summary scripts update the bulk-run and single-cell statistics shown on Hugo and Dash pages. Dash and SequenceServer are deployed as service components with containerized runtime environments, while static Hugo and JBrowse 2 assets are released as web resources. The same directory and configuration structure supports incremental onboarding of additional species. The public repository includes onboarding, data-preparation, and deployment guides, together with configuration templates, species-adapter examples, Docker/Compose files, and ScienceDB demonstration files.

#### ELS prediction

We predicted enhancer-like signatures (ELSs) using matched ATAC-seq and H3K27ac data following an ENCODE-derived strategy (2). The prediction process consisted of two main phases: representative DNA accessibility region (rDAR) identification and ELS scoring/classification.

For rDAR identification, ATAC-seq peaks called by MACS2 were pooled across samples and merged into a non-overlapping representative accessibility-region set using an iterative thresholding and merging procedure. These rDARs were used as candidate accessible regions for ELS scoring.

For ELS scoring and classification, we used matched ATAC-seq and H3K27ac profiles and computed Z-scores from log-transformed, sample-standardized signals. ELSs were defined as rDARs meeting two criteria: (1) ATAC max-Z > 1.64, corresponding to the 95th percentile of a one-sided Gaussian distribution, and (2) H3K27ac max-Z > 1.64. Here, max-Z denotes the maximum standardized signal for a region across the matched sample matrix.

ELSs were classified by genomic distance from transcription start sites (TSS): TSS-proximal ELSs (pELSs) were defined as 0-2000 bp from a TSS, while TSS-distal ELSs (dELSs) were >2000 bp from a TSS. We identified tissue-specific ELSs using Bedtools (35) to analyze non-overlapping ELSs across samples. The main thresholds are described above; implementation-level parameters for rDAR refinement, signal extraction, and ELS classification are provided with the analysis scripts in the GitHub repository.

#### ELS-expression benchmark

We evaluated whether H3K27ac-supported ELSs provided additional information beyond ATAC-only accessibility using the 17 zebrafish benchmark samples. For each sample, we used the sample-specific ELS BED file, ATAC peak BED file, gene annotation table, and RNA expression matrix from the BulkMulti module. Features with midpoints within 2 kb of any TSS were removed so that the analysis focused on distal elements.

ATAC-only peaks were defined as distal ATAC peaks that did not overlap any distal ELS by at least 1 bp. Genes were assigned to groups using a TSS-centered window. The primary analysis used a 25 kb window, and sensitivity analyses repeated the same procedure with 10 kb, 50 kb, and 100 kb windows. A gene was assigned to the ELS-near group if at least one distal ELS fell within the window. Genes without nearby ELSs but with at least one nearby distal ATAC-only peak were assigned to the ATAC-only-near group. Genes without nearby distal ELSs or distal ATAC-only peaks were assigned to the no-ATAC group.

For each sample and window, we compared expression distributions among groups using one-sided Mann-Whitney U tests with the alternative hypothesis that the first group had higher expression. We calculated Cliff’s delta as the non-parametric effect size and adjusted p-values across the 17 samples using the Benjamini-Hochberg procedure. The main comparison was ELS-near versus ATAC-only-near genes; comparisons against the no-ATAC group were used as baseline summaries.

#### ELS-to-gene linkages

We inferred ELS-to-gene linkages using a computational framework adapted from single-cell methodologies (36). The analysis had three steps: (1) ELS identification as described above, (2) generation of sample-specific matrices for ATAC-seq signal at ELSs (sample by ELS matrix) and RNA expression (sample by gene matrix), and (3) calculation of Pearson correlation coefficients between log2-normalized values from both matrices for all potential ELS-gene pairs within a 250 kb window centered on each TSS. We then visualized and explored correlation-based linkages through the miniODP platform.

#### GO and motif analysis

Functional annotation used Gene Ontology (GO) enrichment and motif analysis. We performed GO enrichment with clusterProfiler (version 4.0.5) to identify enriched biological processes and molecular functions. We performed de novo motif discovery and motif enrichment analysis with HOMER (37) to identify TF binding motifs associated with specific genomic regions or biological conditions.

#### Identification of super-enhancers

We identified super-enhancers from H3K27ac profiling data aligned to the GRCz11 reference genome. We used the ROSE algorithm (Rank Ordering of Super-Enhancers) (38) with default parameters to distinguish extended H3K27ac signal domains from typical enhancers.

#### GRN analysis

We constructed GRNs using ANANSE (39), integrating gene expression, ATAC-seq, H3K27ac data, ELS calls, and TF motif annotations. For zebrafish, we built a TF motif database by aggregating vertebrate motifs from the CIS-BP database (Build 2.00) using GimmeMotifs (40). These motifs were converted into a zebrafish-compatible format using the motif2factors function. The same ANANSE binding and network steps were applied to cattle lung and spleen because these samples had matched expression, chromatin, ELS, and motif inputs suitable for tissue-level GRN interpretation. ANANSE estimated TF binding probabilities by combining motif scores with ATAC-seq and H3K27ac signals in ELS regions. Sample-specific GRNs were then constructed by integrating predicted TF binding probability in enhancers, genomic distance between enhancers and target genes, expected TF activity, and expression levels of TFs and target genes. The resulting edges were assigned Network Influence Scores on a 0-1 scale. For visualization and top-TF summaries, edges were sorted by Network Influence Score, the top 100,000 high-scoring edges were retained for each network to keep network display and top-TF extraction consistent, and TFs were ranked by outdegree within this retained subnetwork. The top 10 TFs by outdegree were used for the GRN summaries. We visualized network plots using the igraph R package (version 1.3.1).

#### Literature support curation for top TFs

We curated literature support for the top 10 TFs from four GRNs: zebrafish blood, zebrafish brain, cattle lung, and cattle spleen. We collected supporting evidence from manuscript-cited references and PubMed searches and classified each TF as direct, indirect, or unsupported. Direct support required abstract-level evidence that the TF has a function in the corresponding tissue, cell type, or core biological process, with species match noted when available. Indirect support included evidence from related species, related TF family members, disease settings, or related tissue processes. Unsupported TFs lacked sufficient evidence under these criteria. The curated evidence, PMID list, and classification for all 40 TFs are provided in Supplementary Table 4.

#### Resource comparison

We compared miniODP with selected species-focused omics resources using public resource descriptions, associated papers, and miniODP implementation documentation. The comparison focused on resource type, scale, data source, main offering, reusable deployment, new-species onboarding, and processing workflow. Reusable deployment was scored as present when public evidence supported redeployment of a similar portal architecture through code, containers, or deployment documentation. New-species onboarding was scored as present when a resource provided a documented process for adding a new organism to an existing instance; resources that described future expansion or data submission without such a documented workflow were marked as not documented, and single-species resources were marked as single species. Processing workflow was scored as present or partial when a resource provided reusable data-processing workflows, methods-level processing details, notebooks, or public scripts for its main data objects. Partial indicates component-level evidence that is insufficient for full third-party redeployment, onboarding, or process reproduction. Table entries use ND for not documented, SS for single species, and NA for not applicable. Size values were treated as scale indicators and were not used as ranking criteria. The evidence checked for each resource is summarized in Supplementary Table 5.

#### Code and demonstration files

The analysis and portal resources are maintained with documented code, configuration files, and example data. The analysis package includes source code for preprocessing, quality control, ELS prediction, ELS-to-gene linkage, GRN construction, and downstream summaries. The portal implementation includes module-specific configuration files, species display settings, species adapters, data-conversion scripts, onboarding and deployment documentation, Docker/Compose files, and demonstration files for data analysis and visualization. Public links are provided in Data Availability.

## Results

### miniODP framework design

We designed miniODP (https://tulab.genetics.ac.cn/miniodp/) as a reusable framework for building multi-omics resources in understudied organisms (Figure 1A). Here, reusable means that public code, configuration templates, species adapters, data-conversion scripts, deployment documentation, and demonstration files support portal reuse and new-species onboarding. The architecture is configurable and can be extended to new organisms without rebuilding the portal for each species.

**Figure 1:**
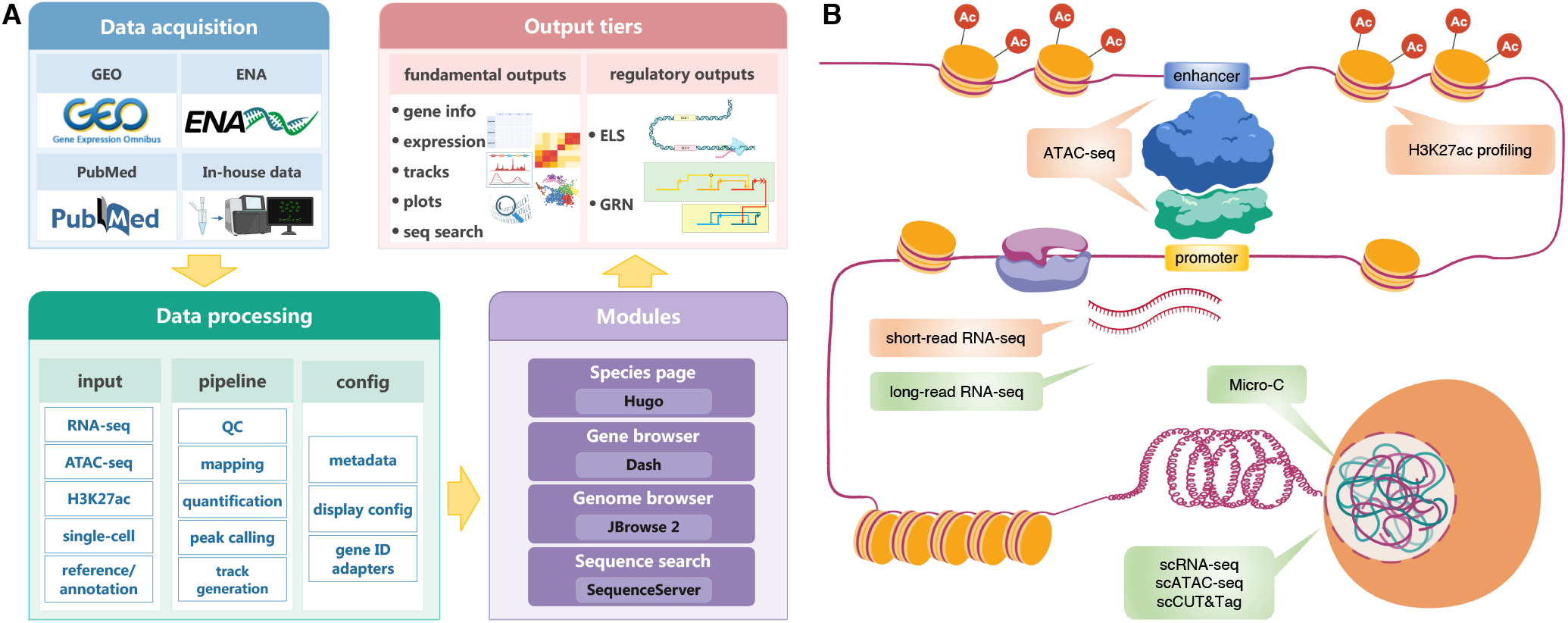
Overview of the miniODP framework and miniENCODE core assay design. (A) miniODP portal framework linking data acquisition, data processing, web modules, and tiered outputs. General outputs support gene information, expression views, genome tracks, plots, and sequence search, whereas regulatory outputs such as ELSs and GRNs are generated when suitable matched data and analysis contexts are available. (B) miniENCODE core assay design, with RNA-seq, ATAC-seq, and H3K27ac profiling as the core layers and long-read RNA-seq, Micro-C, and single-cell assays as optional extensions. The illustration was partially generated using biogdp.com (41).

The public site is organized under a unified entry point and combines four connected components: Hugo species pages (https://gohugo.io/), gene-centric interactive applications built with the Python Dash framework (https://dash.plotly.com/), JBrowse 2 genome browser sessions (30), and SequenceServer-based sequence search (31) (Figure 2). Species pages provide the entry for each organism and summarize available bulk and single-cell resources. Dash applications provide configuration-driven gene search and annotation, quantitative expression views, single-cell views, and scATAC gene-activity views; when matched core-assay data are available, BulkMulti panels also combine expression, ATAC signal, ELS coordinates, and ELS-to-gene linkages. Depending on available annotations, searchable fields can include Ensembl-style gene IDs, gene names, GO terms, InterPro domains, TF families, and orthologs; gene pages can also include Ensembl links (32), TF-family annotations linked to AnimalTFDB (33), and InterPro domain annotations (34). JBrowse 2 provides genome-context visualization of omics tracks, and SequenceServer supports searches against genome, cDNA, and protein databases.

**Figure 2:**
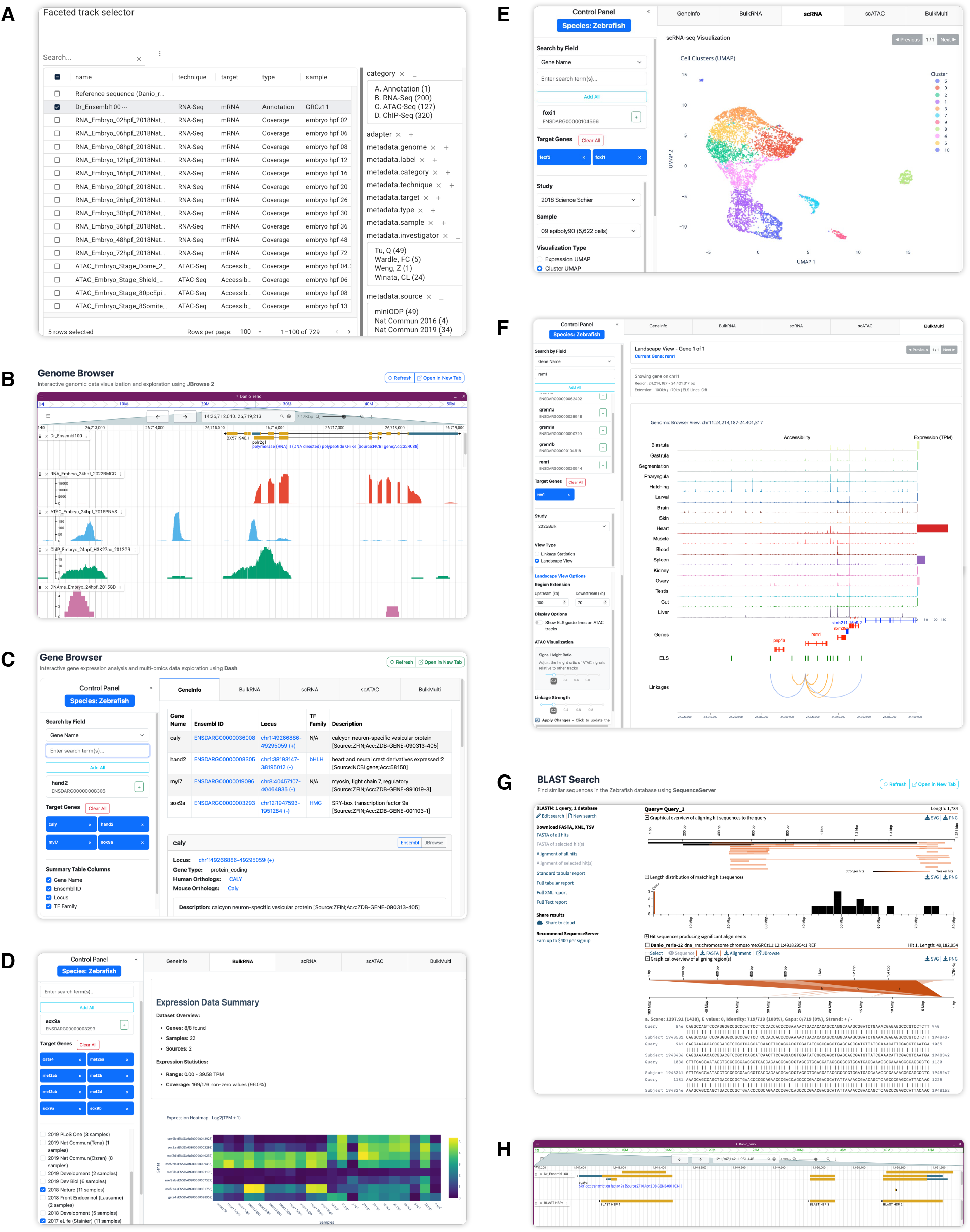
Functional modules of miniODP. (A, B) The genome browser supports locus-level visualization of multi-omics tracks and track selection by sample, assay, data type, target molecule, and data source. (C–F) The gene browser links gene annotations, expression summaries, single-cell views, and, where matched core-assay inputs are available, multi-omics panels including ELS-gene linkages. (G, H) SequenceServer provides BLAST search against genome, cDNA, and protein databases, with hits linked back to genome-browser coordinates.

The modules are connected around gene- and locus-level navigation. Users can start from a species page, open genome-browser views for the corresponding assembly, move to Dash gene pages for configured gene-level searches and quantitative views, and use SequenceServer for genome, cDNA, or protein queries. Gene-search fields include gene identifiers and gene names for all species, with GO terms, InterPro domains, TF families, and orthologs added where the corresponding annotations are available. JBrowse 2 track selectors use experimental category, sample type, data source, target molecule, technique, and data type, so large track sets can be filtered without manually scanning all files. Gene pages and SequenceServer results can also link back to genome-browser coordinates, allowing genes and sequence matches to be interpreted in their genomic and regulatory context. Together, these links make the portal a connected workflow rather than a set of independent viewers.

This framework separates species configuration from application logic. Species-specific public information, external database links, summary statistics and data directories are all managed by configuration files. Gene-centric retrieval is processed by species adapters, which support both standard and local gene identifiers, including species that use dual identifier systems. Therefore, species-specific handling is restricted in configuration files and adapters, instead of being coded in the application logic. The same Dash code can process species with different annotation rules.

A set of data processing pipelines is provided. The conversion scripts are used to prepare gene information, bulk RNA-seq tables, single-cell data objects, JBrowse references and tracks, and SequenceServer database manifests. The summary and maintenance scripts are used to update the statistics of bulk runs and single-cell datasets in Hugo and Dash. This layer connects upstream data curation with portal release and maintenance.

Deployment is organized modularly. Static pages and genome browser resources are synced as web resources, and Dash and SequenceServer run in containerized services. Data directories and application code are mounted separately. Therefore, a new species can start with a minimal dataset, including genome, gene information table, and some bulk expression tables. More genome browser tracks, BLAST databases, single-cell modules and multi-omics outputs can be added when data are available.

This tiered design is for a practical limit: the completeness of data types in non-model organisms varies greatly. Species with matched core assays support extra ELS-centric analysis and even GRN construction with enough data. Species with partial data can still provide gene-and genome-centric functions like browsing, search, and visualization. In the current instance, miniODP uses this structure for all seven species, supporting different levels of downstream analysis with a common architecture.

### Resource-efficient core assay design

For communities that cannot produce ENCODE-scale data, we propose a small set of omics profiling assays. The complete ENCODE toolbox covers DNA-protein interactions, transcription, chromatin accessibility, DNA methylation, RNA binding, and 3D chromatin architecture (2). This broad coverage is useful for systems with enough resources. But most non-model organisms cannot replicate such scale. The complete toolbox requires high cost, many samples, specialized protocols, and infrastructure for data processing and presentation. For miniODP, the practical goal is to select a small set of assays to support the most fundamental regulatory analysis.

We term this set of three assays miniENCODE: short-read RNA-seq, ATAC-seq and H3K27ac profiling (Figure 1B). The purpose of this set is to provide a starting point for resource-limited projects, not an ENCODE-equivalent replacement. We chose them because they are widely used, applicable in multiple species, and directly link to the portal outputs. Together they provide transcriptional state, chromatin accessibility and active regulatory marking, which are the minimal requirements for ELS and GRN analysis.

RNA-seq is the anchor as it provides the expression matrix, which is used for sample comparison, gene-level interpretation, ELS-to-gene linkage and GRN construction. ATAC-seq maps accessible regions, which are used to define candidate chromatin space. H3K27ac restricts this space further to active enhancer-like and promoter-associated states (2). In the zebrafish benchmark, H3K27ac profiles were generated by CUT&Tag. Compared with traditional ChIP-seq, CUT&Tag needs less sample input and sequencing, and produces higher signal-to-noise in histone-mark profiling (20). Therefore, CUT&Tag is suitable for projects with limited material and sequencing budget.

These three assays directly link to the outputs of miniODP. ATAC-seq and H3K27ac together define enhancer-like signatures (ELSs), while RNA-seq provides expression measurements for gene-level interpretation, ELS-to-gene linkage and GRN construction. The reason to include H3K27ac is that open chromatin alone cannot distinguish active regulatory regions from other accessible loci. We did not include H3K4me3, because it mainly resolves promoter activity, while H3K27ac marks active enhancers and promoters simultaneously, providing wider coverage at the same assay budget. Together, these core assays define the minimal dataset required for regulatory elements and network analysis, but are not intended to cover all functional layers ENCODE provides.

When resources permit and investigative questions require, more assays could be used. Long-read RNA-seq could improve transcript models, which is particularly useful for newly sequenced genomes when transcript boundaries and isoforms are uncertain (42). Micro-C or related chromosome-conformation assays could provide long-range contact information, and could be used to interpret distal regulatory elements (43, 44). Single-cell RNA-seq, single-cell ATAC-seq or single-cell CUT&Tag could resolve cell-type-specific expression and regulatory states in complex tissues (45–48). Spatial assays and additional histone marks can serve as optional layers.

### Zebrafish benchmark: data coverage

We used zebrafish as the deep benchmark because it combines experimental tractability with broad public omics coverage. We collected and curated zebrafish datasets from GEO (49) and ENA (50), then organized them in miniODP as a data type matrix by biosample and assay type (Figure 3). At the portal level, this matrix includes RNA-seq, ATAC-seq, ChIP-seq/CUT&Tag, DNA methylation, scRNA-seq, and scATAC-seq across eight developmental stages and 20 embryonic and adult tissues. For ChIP-seq and CUT&Tag datasets, tracks were further grouped by target type, with H3K27ac treated as the core histone-mark layer for the assay design.

**Figure 3:**
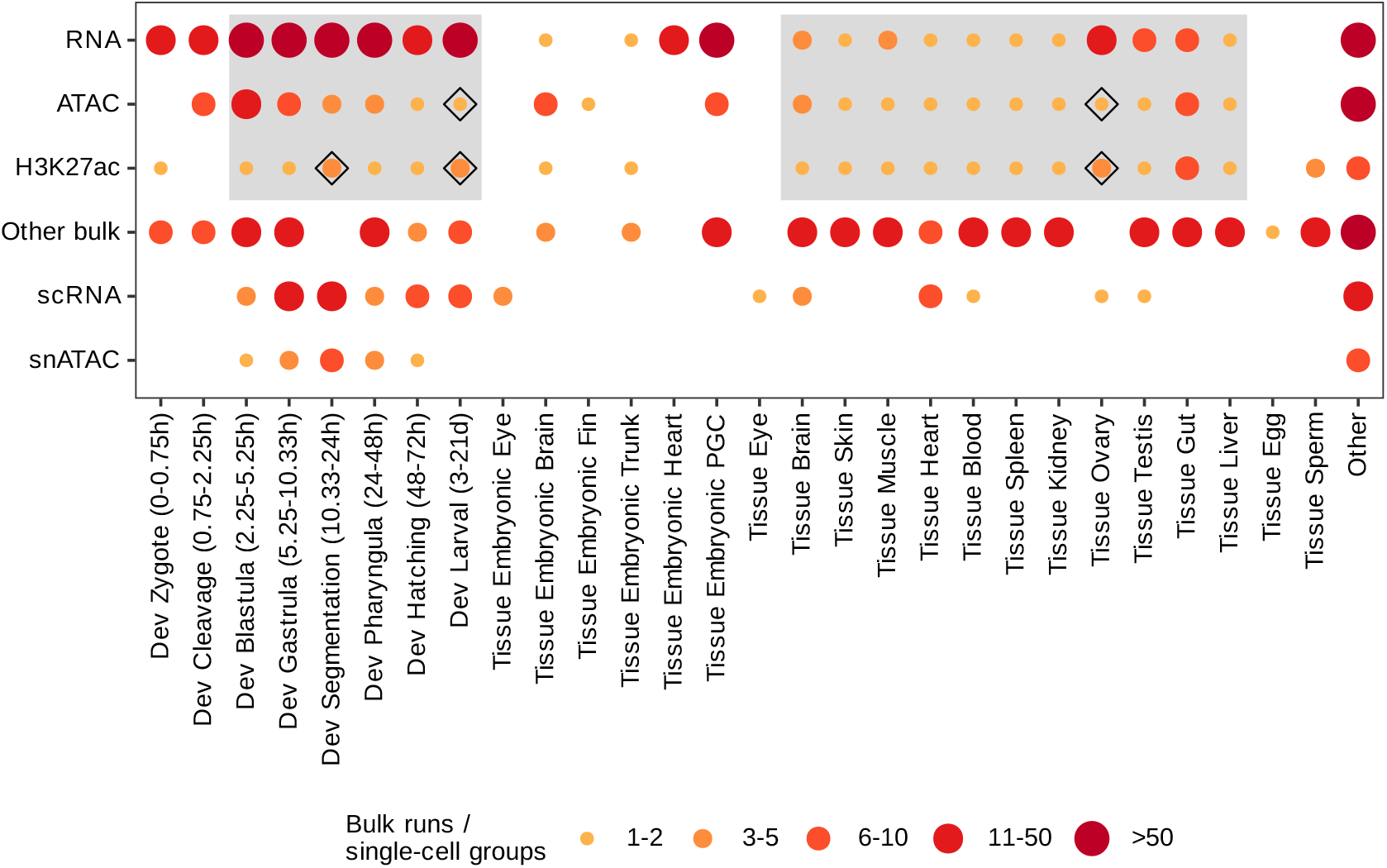
Zebrafish data coverage used for the benchmark analyses. The data type matrix summarizes omics datasets by biosample and assay type. Dot size and color intensity indicate the number of bulk runs for bulk-assay rows and the number of single-cell sample groups for single-cell rows. Bulk runs denote run-level bulk omics sequencing records, whereas single-cell sample groups denote curated matrix-level display units rather than cell clusters. Grey boxes highlight the core miniENCODE assay targets (RNA-seq, ATAC-seq, and H3K27ac profiling) for developmental stages and adult tissues. Other bulk includes histone-mark ChIP/CUT&Tag beyond H3K27ac, TF ChIP/CUT&Tag, DNA methylation, and other bulk assays. Diamonds indicate datasets generated in this study, with quality-control summaries shown in Supplementary Figures 2 and 3.

For the zebrafish benchmark, we first considered the coverage of the three core miniENCODE assays across samples. These samples included six developmental stages from blastula to larval stage, and 11 adult tissues: brain, skin, muscle, heart, blood, spleen, kidney, ovary, testis, gut, and liver. Public datasets already covered most of the matrix, but lacked matched ATAC-seq and H3K27ac profiles for a few samples. Therefore, we generated ATAC-seq datasets for the larval stage and ovary tissue, and H3K27ac CUT&Tag datasets for the segmentation stage, larval stage, and ovary tissue. These newly generated datasets are marked by diamonds in Figure 3. Quality-control analyses of these libraries showed genomic feature enrichment, high chromosome-mapping rates, and reproducibility between biological replicates (Supplementary Figures 2 and 3). Together with public datasets, these generated datasets formed a 17-biosample benchmark matrix, in which each sample had matched RNA-seq, ATAC-seq, and H3K27ac inputs. This matrix was used for ELS identification, ELS-expression benchmarking, ELS-to-gene linkage, and GRN construction.

Each dataset was manually curated for experimental design, biological replicates, sequencing depth, and available quality metrics. The selection criteria prioritized replicate support, metadata completeness, and coverage of key developmental stages or tissues, rather than inclusion of all available public tracks.

The final data type matrix is shown in Figure 3. The plot compares coverage across assays, developmental stages, and tissues. It also separates histone-modification tracks and TF-oriented ChIP/CUT&Tag datasets, and uses H3K27ac as the core histone mark in the assay design. The matrix shows both the breadth of zebrafish public data and the subset of core assays needed for downstream regulatory analyses.

### Zebrafish benchmark: enhancer-like signatures

Enhancer activity is commonly associated with open chromatin, H3K27ac, and binding of cell type-specific TFs (51, 52). ATAC-seq profiles accessible chromatin genome-wide (53), while H3K27ac marks active enhancer and promoter states (54). Using the 17-sample three-assay matrix, we applied an ENCODE-derived strategy to predict enhancer-like signatures (ELSs) (2). This pipeline first identified representative DNA accessibility regions (rDARs) from ATAC-seq peaks, then calculated Z-scores of ATAC-seq and H3K27ac signals. We defined rDARs with both ATAC-seq and H3K27ac signals above the 95th percentile (Z-score ≥ 1.64) as ELSs, and divided them into TSS-proximal ELSs (pELS, 0–2 kb from a TSS) or TSS-distal ELSs (dELS, >2 kb) according to the distance from transcription start sites (TSS).

This procedure identified 52,350 ELSs, representing 1.2% of the zebrafish genome and 17% of ATAC-seq peaks; 37% were pELSs and 63% were dELSs. Compared with previous zebrafish regulatory-element resources (7, 9), 20% of the ELSs were not overlapped by those annotations (Supplementary Figure 4). All ELSs are available through the miniODP genome browser, where ELS calls can be viewed together with RNA-seq, ATAC-seq, and H3K27ac tracks in representative tissues such as heart, brain, intestine, and liver (Figure 4A).

**Figure 4:**
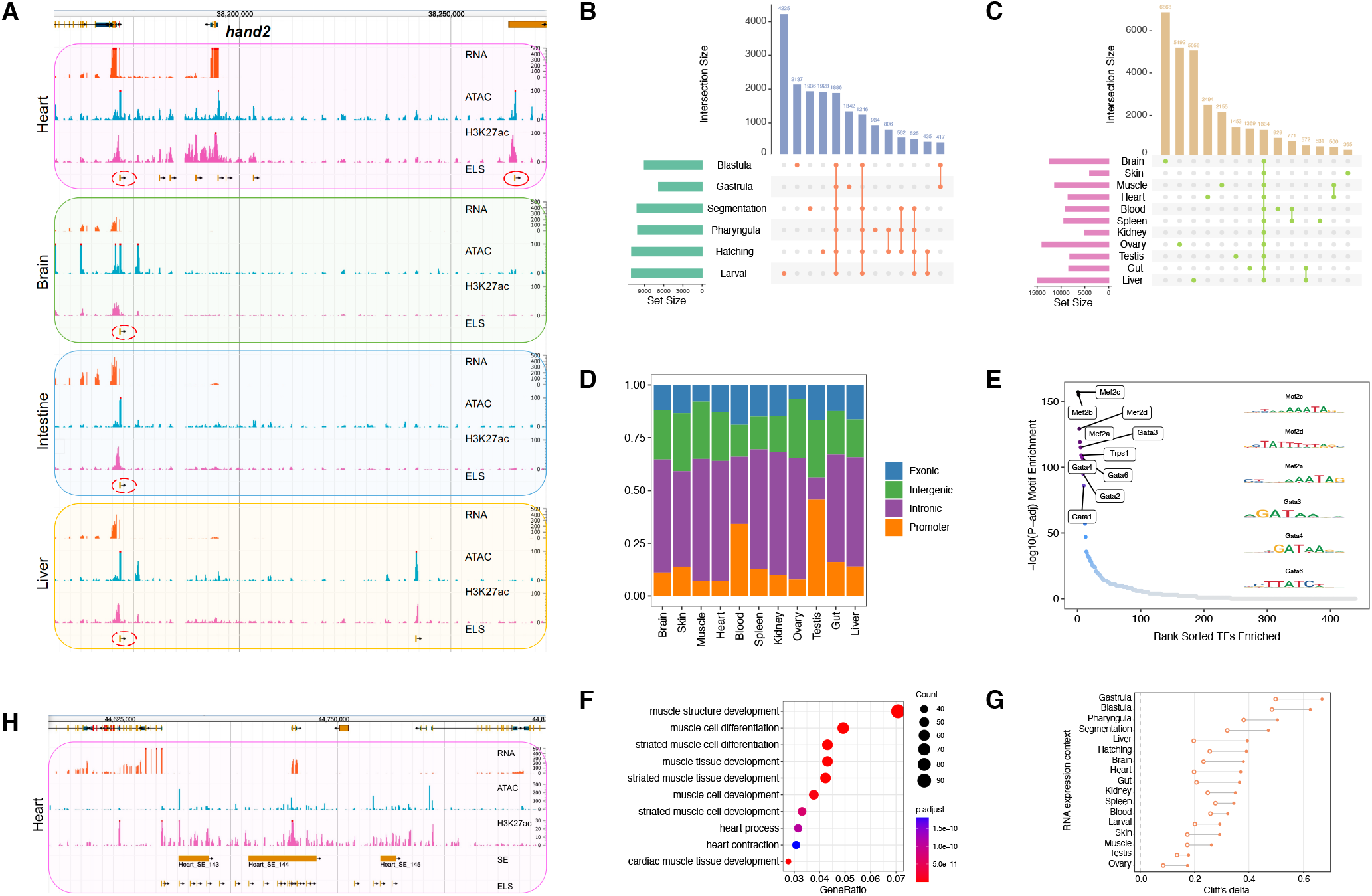
Zebrafish enhancer-like signatures and associated annotations. (A) Multi-omics visualization of ELSs in four representative tissues (heart, brain, intestine, and liver). The genomic view displays gene annotations followed by tissue-specific profiles. Each tissue panel shows RNA-seq, ATAC-seq, H3K27ac, and predicted ELS tracks. ELS identification requires co-occurrence of ATAC-seq and H3K27ac signals. The dashed left ellipse highlights a ubiquitously active ELS, and the solid right ellipse shows heart-specific ELS22 selected for reporter testing. (B, C) Upset plots illustrating ELS activation patterns across developmental stages (B) and tissues (C). Horizontal bars represent individual stages or tissues, and vertical bars show intersection sizes. The larval stage contains the highest number of stage-specific ELSs, while brain tissue shows the most tissue-specific ELSs. We identified 1,886 developmentally persistent and 1,334 tissue-ubiquitous ELSs. (D) Genomic distribution of tissue-specific ELSs, showing predominant localization in intronic (51%) and intergenic (23%) regions. (E) TF binding motif enrichment analysis for heart-specific ELSs. TF motifs are ranked by enrichment significance, with motifs of six cardiac regulators highlighted. (F) Gene Ontology analysis of genes associated with heart-specific ELSs, highlighting heart- and muscle-related terms. (G) ELS-expression benchmark at a 25 kb TSS window. Filled circles show Cliff’s delta for ELS-near genes versus no-ATAC genes, open circles show ATAC-only-near genes versus no-ATAC genes, and grey lines connect the two comparisons within each RNA expression sample. The primary comparison is ELS-near versus ATAC-only-near genes, with no-ATAC genes shown as a baseline. (H) Genomic architecture of super-enhancers, characterized by extended regions with strong ATAC-seq and H3K27ac signals.

The ELS atlas captured both shared and sample-restricted regulatory activity. Among the 52,350 ELSs, 51% (26,695) showed stage- or tissue-specific activity. Figure 4A shows a ubiquitously active ELS and the heart-specific ELS22, which had strong ATAC-seq and H3K27ac signals in heart samples and was selected for reporter testing below. The larval stage contained the highest number of stage-specific ELSs, while brain, ovary, and liver contained the largest numbers of tissue-specific ELSs (Figure 4B,C). We also identified 1,886 developmentally persistent, 1,334 tissue-ubiquitous, and 702 universally active ELSs (Figure 4B,C; Supplementary Table 1). These categories separate elements active in restricted samples from elements active across many samples. Stage-specific and tissue-specific ELSs provide candidate sets for transgenesis. Constitutively active ELSs provide candidate loci for knock-in designs, including cases where random *Tol2* integration is not desirable because transgene expression can be affected by local chromatin context (55).

**Table 1.**
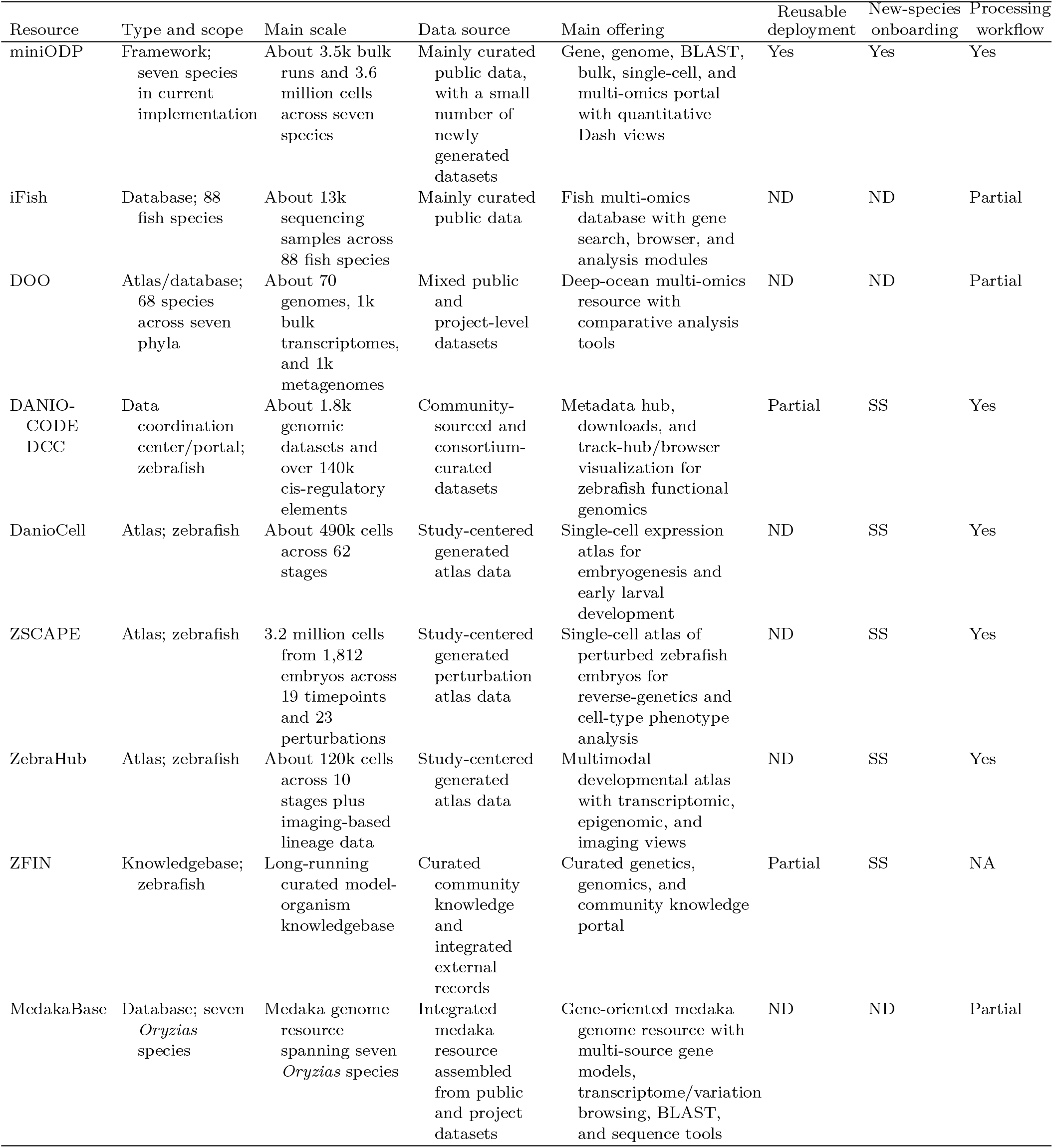
Comparison of miniODP with selected species-focused omics resources. Size is shown as a scale indicator and is not used as a ranking criterion. Reusable deployment, new-species onboarding, and processing workflow were scored from public resource descriptions, associated papers, and documentation; scoring definitions are provided in Methods, and the evidence checked for each resource is summarized in Supplementary Table 5. ND, not documented; SS, single species; NA, not applicable.

The overlap structure differed between developmental stages and adult tissues. Developmental stages showed substantial ELS sharing, whereas distinct adult tissues showed more sample-restricted ELS activity (Figure 4B,C). Shared ELSs were most apparent in tissue groups with related developmental origins or functions, including muscle and heart, blood and spleen, and gut and liver (56, 57). This pattern is consistent with the expectation that regulatory activity is more similar between related tissues or temporally adjacent stages than between unrelated adult organs. Genomic localization analysis showed that tissue-specific ELSs were mainly intronic (51%) or intergenic (23%) (Figure 4D and Supplementary Figure 5), similar to patterns reported in mouse enhancers (58). Testis-specific ELSs showed a different distribution, with a lower intronic fraction and higher promoter-region enrichment than other tissues. This observation was kept as a descriptive feature of the atlas because the current dataset was not designed to resolve the mechanism of testis-specific promoter-proximal activity.

We next examined whether tissue-specific ELSs were consistent with tissue annotations. Motif enrichment was performed for tissue-specific ELSs, and TF motifs were ranked by enrichment significance. In heart-specific ELSs, six cardiac regulators appeared among the top ten enriched TF motifs: Mef2c, Mef2d, Mef2a, Gata3, Gata6, and Gata4 (Figure 4E). Similar tissue-relevant motif enrichments were observed in other tissues (Supplementary Figure 6). GO analysis of genes adjacent to tissue-specific ELSs also recovered tissue-relevant terms: heart-specific ELS-associated genes were enriched for heart contraction and muscle development, brain-specific ELSs for neuronal differentiation and axon development, liver-specific ELSs for metabolic processes, and testis-specific ELSs for cilium organization and cell-cycle regulation (Figure 4F and Supplementary Figure 7). These annotations indicate that tissue-specific ELSs are often located near genes with functions matching the corresponding sample.

We then tested whether H3K27ac-supported ELSs were more closely associated with active genes than distal ATAC-only peaks in this benchmark. For each of the 17 zebrafish samples, we assigned genes to three groups based on nearby distal features: ELS-near, ATAC-only-near, and no-ATAC. At a 25 kb TSS window, ELS-near genes showed higher expression distributions than ATAC-only-near genes across samples, reflected by consistently higher ELS-near effect sizes relative to the no-ATAC baseline (Figure 4G; Supplementary Figure 8). The effect sizes were modest in magnitude, but the direction was stable. This supports using H3K27ac together with ATAC-seq for ELS definition in miniODP.

We also generated super-enhancer (SE) annotations from H3K27ac profiles using ROSE (38). SEs were treated as extended H3K27ac-marked regions, following the original ROSE definition of stitched enhancer domains (38, 59). In the heart example, H3K27ac-density ranking separated predicted SEs from typical enhancers, and GO analysis of SE-associated genes recovered heart morphogenesis, vasculogenesis, and cardiocyte differentiation terms (Supplementary Figure 9A,B). Across embryonic stages and adult tissues, we identified hundreds of SEs per sample (Supplementary Figure 10A). These regions spanned larger domains with strong ATAC-seq and H3K27ac signals (Figure 4H). SEs had a median length of 28,344 bp, compared with 297 bp for individual ELSs, and each SE contained an average of two ELSs; 86% contained at least one ELS (Supplementary Figure 10B,C). These SE annotations add a broader regulatory track layer for inspecting extended H3K27ac domains around genes of interest. They can be browsed together with ELS, ATAC-seq, H3K27ac, and RNA-seq tracks, and are intended for exploratory interpretation rather than as the primary benchmark for the core assay design.

### Zebrafish benchmark: selected reporter assays

We used transient zebrafish reporter assays to test whether selected heart-associated ELSs could drive detectable reporter activity in vivo. From the 2,494 heart-specific subset of 8,525 heart-activated ELSs, we selected 23 candidates for GFP reporter testing (Supplementary Table 2). Candidate sequences were cloned into the pSTG vector upstream of a minimal EF1a promoter-EGFP cassette and injected into one-cell-stage zebrafish embryos with transposase mRNA. Embryos were examined at 3 dpf, and at least 100 embryos were scored for each construct.

These assays were performed as F0 transient reporter tests, so mosaic expression was expected. We therefore interpreted the results as selected-candidate support, with stable-line experiments required for stronger claims about spatial specificity. Using a 5% GFP-positive embryo threshold, 15 of 23 tested ELSs (65%) exceeded the threshold for detectable reporter activity (Figure 5A). Among these positive constructs, GFP signals were predominantly observed in the heart, with additional signals in muscle and eye regions for some constructs (Figure 5B,C). Wider-field embryo images are provided in Supplementary Figure 11. The eight candidates without detectable reporter activity may reflect temporal differences between the omics samples and the 3 dpf assay time point, missing sequence context in the cloned fragments, or false-positive ELS predictions.

**Figure 5:**
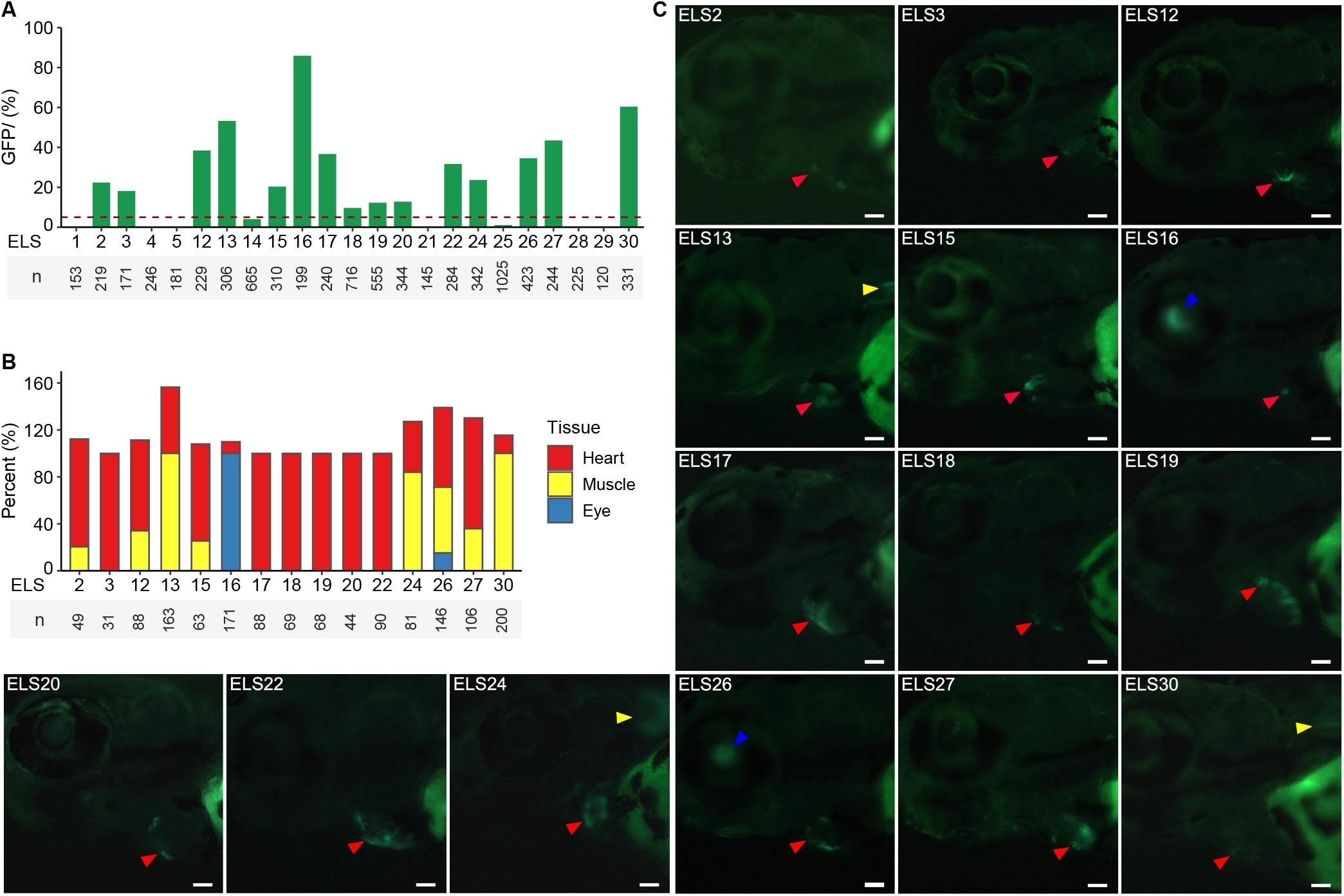
Selected F0 reporter assays for heart-associated ELSs in zebrafish. (A) Percentage of GFP-positive embryos for each tested ELS construct. The dashed line indicates the 5% threshold used to define detectable reporter activity; 15 of 23 constructs exceeded this threshold. (B) Tissue signal annotations for constructs with detectable reporter activity. Tissue categories are not mutually exclusive because individual constructs can show GFP signals in more than one tissue; therefore, summed percentages can exceed 100%. (C) Representative GFP patterns in F0 transient embryos. Red, yellow, and blue arrowheads indicate heart, muscle, and eye signals, respectively, as summarized in panel B. Scale bar, 50 µm.

ELS22 provides one example linking the reporter assay to the ELS predictions. This element showed heart-enriched ATAC-seq and H3K27ac signals and drove detectable GFP expression in the heart in the transient assay. These assays provide candidate-level support for a subset of heart-associated ELSs. Functional testing of additional elements or stable transgenic lines would be required to establish spatial activity for individual ELSs in greater detail.

### Zebrafish benchmark: gene regulatory networks

We next linked ELS activity to gene expression and used matched ELS, chromatin, motif, and expression information to construct sample-level gene regulatory networks (GRNs). For ELS-to-gene linkage, we adapted a correlation-based strategy from single-cell regulatory analysis (36). ELS accessibility profiles were derived from ATAC-seq signal matrices, and gene expression profiles were derived from RNA-seq matrices across the 17 zebrafish benchmark samples. For candidate ELS-gene pairs within a 250 kb window centered on each TSS, we calculated correlation-based linkage scores ranging from −1 to 1. This linked the 52,350 ELSs with 26,354 genes and produced the linkage tracks shown in the miniODP multi-omics view (Figure 6A).

**Figure 6:**
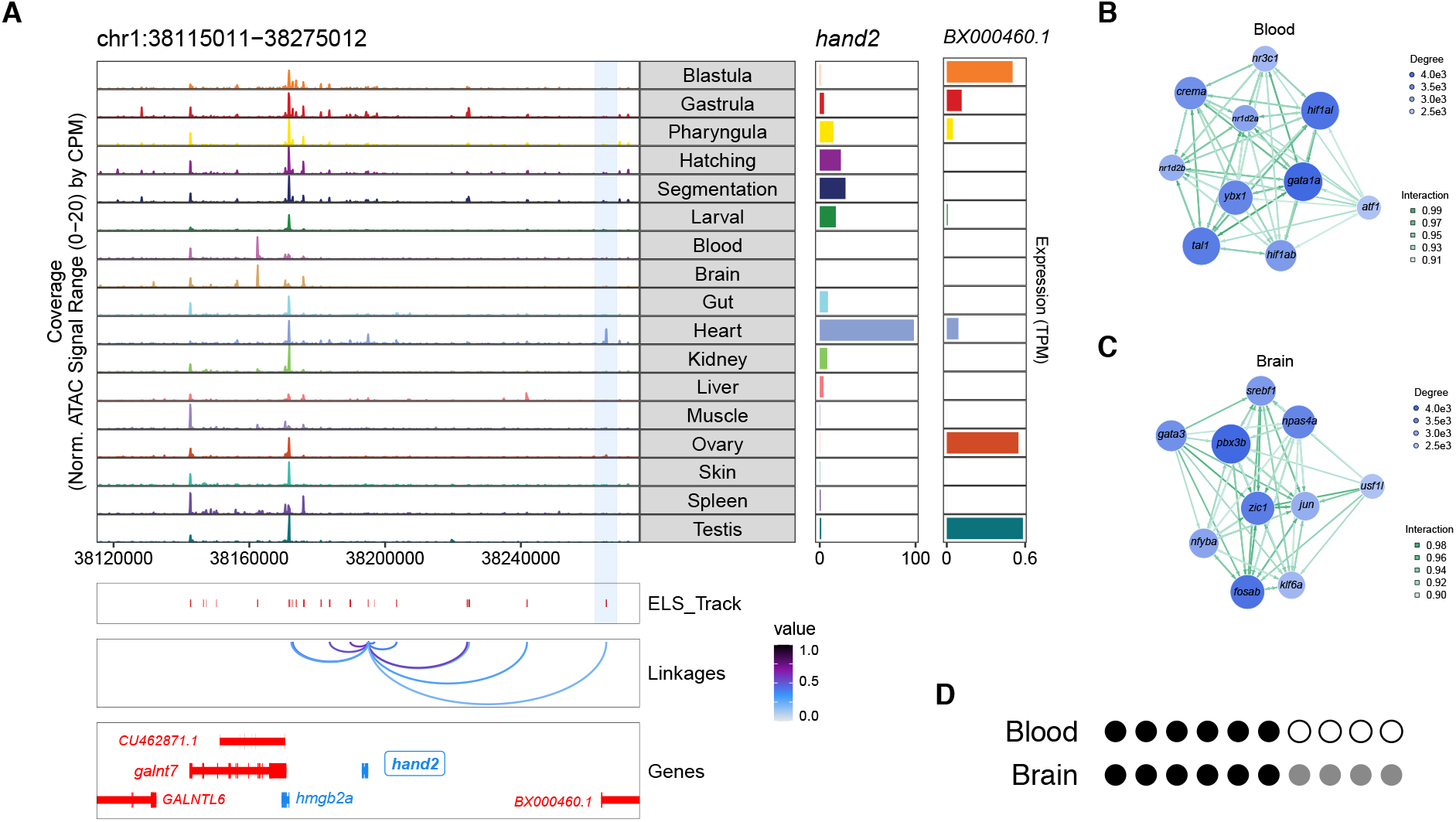
Zebrafish ELS-gene linkages and GRN outputs. (A) Multi-omics visualization of *hand2* regulatory elements. The central panel shows ATAC-seq tracks across samples, the right panel displays RNA-seq-derived gene expression levels, and the bottom panel depicts predicted ELSs, ELS-gene linkages, and associated genes. *hand2* shows heart-specific expression, with ELS22 located approximately 70 kb downstream and showing heart-enriched chromatin accessibility and H3K27ac signal. (B, C) Tissue-specific enhancer-gene regulatory sub-networks for blood (B) and brain (C). Blue circles denote the top 10 TFs in each tissue, ranked by outdegree within the top 100,000 edges. Circle size and color intensity reflect the number of predicted target genes connected to each TF, and edge color intensity indicates interaction strength. These networks provide candidate-prioritization outputs. (D) Literature-support summary for the top 10 TFs in the blood and brain GRNs. Each circle represents one TF among the top 10; circles are grouped by evidence level. Filled black circles indicate direct tissue-relevant support, filled gray circles indicate indirect support, and open circles indicate unsupported candidates under the current curation criteria. Detailed evidence is provided in Supplementary Table 4.

The *hand2* locus illustrates how these linkages are used in the portal. *hand2* is a cardiac developmental regulator (60) and showed heart-specific expression in our dataset. ELS22, located approximately 70 kb downstream, showed heart-enriched accessibility and H3K27ac signal. ELS22 was linked to *hand2* by positive correlation between chromatin accessibility and gene expression (Figure 6A). The same ELS lies within an intron of *BX000460*, which showed weak expression in ovary and testis but not in heart. This example shows why the framework uses sample-matched chromatin and expression profiles instead of assigning distal elements only by nearest gene or host-gene location.

Building on these ELS-gene linkages, we constructed zebrafish GRNs using ANANSE (39). We first built a zebrafish TF motif database using GimmeMotifs (40). ANANSE then estimated TF binding probabilities by combining motif scores with ATAC-seq and H3K27ac signals in ELS regions. TF-gene binding scores were calculated by integrating enhancer intensity and motif evidence within 100 kb of each TSS, and these scores were combined with TF activity and expression levels of TFs and target genes to generate interaction scores on a 0–1 scale. The resulting networks provide regulator-prioritization outputs for each sample; functional testing is required to evaluate individual TF predictions. We examined the top 10 TFs in the blood and brain GRNs, and curated literature support for each TF (Figure 6B–D and Supplementary Table 4). In the blood GRN, six TFs had direct evidence in zebrafish hematopoiesis, including *hif1al, hif1ab, gata1a, tal1, ybx1* and *nr3c1*. For example, Hif-1*α*/Hif-2*α* regulate hemogenic endothelium and HSC formation (61), and *nr3c1*-encoded glucocorticoid receptor signaling controls embryonic HSPC production (62). We did not find tissue-matched evidence for the other four candidate TFs. In the brain GRN, six TFs had direct zebrafish neural evidence, including *zic1, npas4a, fosab, gata3, klf6a* and *jun*; the other four had indirect support from related systems or TF families. For instance, the AP-1 component *jun* is linked to zebrafish brain development through the Trim69/AP-1 pathway (63).

If only expression level is considered, some high-ranked TFs are not tissue-specific genes. For example, *hif1ab* and *nr3c1* are expressed in a wide range of tissues, but have stronger connectivity in the blood GRN; *jun* is also widely expressed, but has stronger connections in the brain GRN (Figure 6B,C and Supplementary Figure 12). This indicates that GRN predictions integrate more layers of information than expression, and the GRN layer could be used for candidate prioritization.

### Seven-species implementation and tiered outputs

Next, we applied the same portal architecture to other species. Currently, miniODP contains seven species: zebrafish, medaka, turquoise killifish, Mexican tetra, Hydra, lancelet and cattle. The portal has curated 3,568 bulk runs, 124.5 billion reads, 155 single-cell sample groups, 3.61 million cells and 1,865 genome-browser tracks. The data distribution is uneven, as expected for non-model and emerging model organisms: zebrafish contributes the largest single-cell component; medaka and killifish contribute large bulk RNA-seq collections; cattle, Hydra, lancelet and Mexican tetra provide more focused matched regulatory datasets drawn from a few studies.

This implementation separates two levels of output. When matched RNA-seq, ATAC-seq, and H3K27ac data were available for a sample, miniODP produced ELS-centered outputs including ELSs, tissue- or stage-specific ELS summaries, motif enrichment, and GO enrichment. This applied to zebrafish and to four additional species with focused three-assay contexts: cattle, Hydra, lancelet, and Mexican tetra (Figure 7A). GRN construction was applied more narrowly to samples with suitable matched expression, chromatin, and TF motif inputs; in the current multi-species implementation, these examples were zebrafish benchmark samples and cattle lung and spleen. Medaka and killifish were handled as partial-data cases. Their H3K27ac coverage is insufficient for systematic ELS-centered analysis or GRN construction under the core assay definition, but both can still be served through the core portal modules.

**Figure 7:**
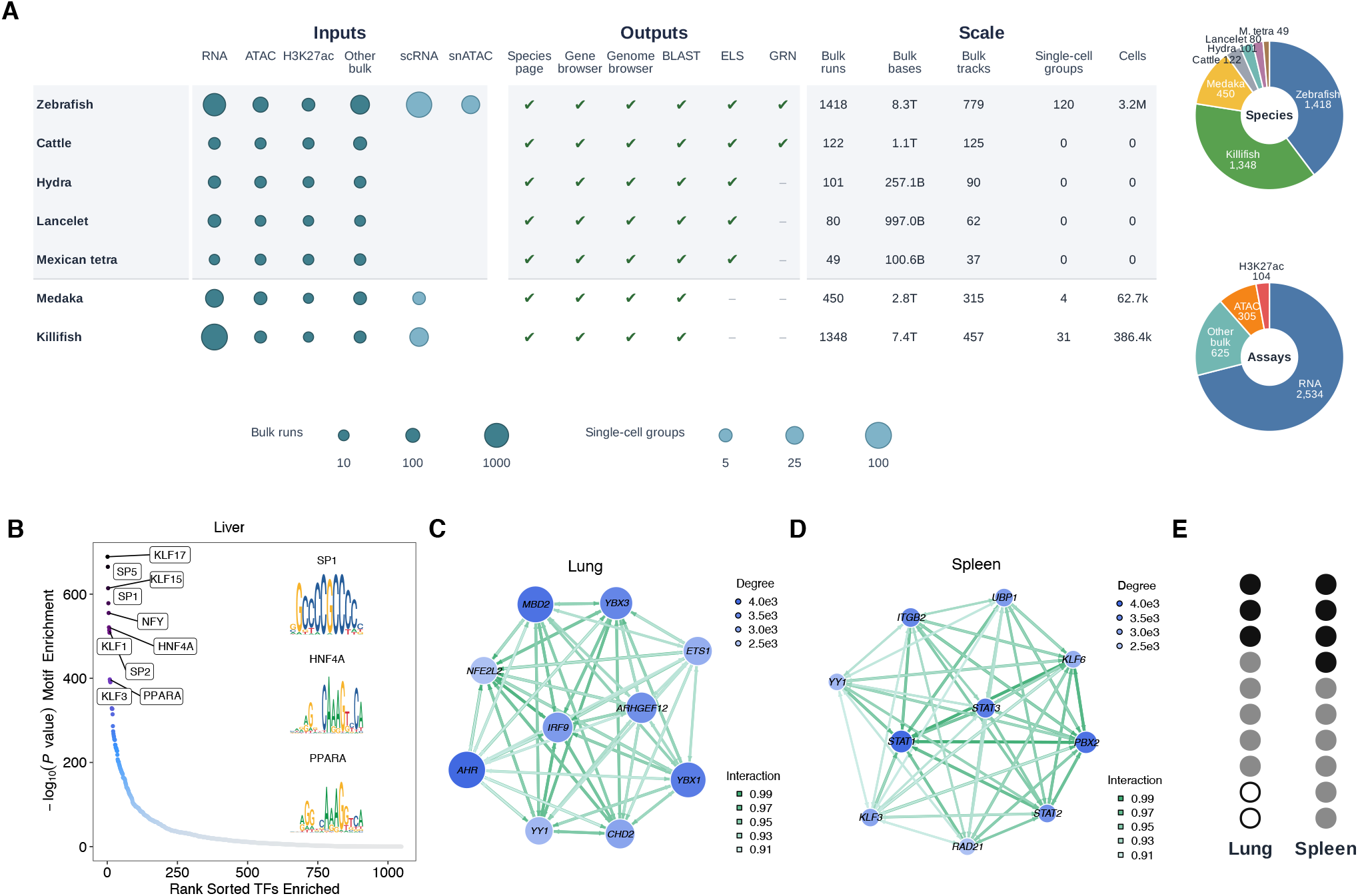
Seven-species implementation and matched regulatory outputs in miniODP. (A) Species-by-assay and output matrix for the seven-species portal. Dot size represents the number of bulk runs or curated single-cell sample groups for each species and data type, using the same definitions as Figure 3. Checkmarks indicate available portal or analysis outputs. Donut charts summarize bulk-run composition by species and assay class. (B) Ranked list of TF motifs enriched in cattle liver ELSs. TFs are sorted by motif-enrichment significance. (C, D) Enhancer-gene regulatory sub-networks for cattle lung (C) and spleen (D). Blue circles represent the top 10 TFs in each tissue, ranked by outdegree within the top 100,000 edges. Circle size and color intensity correspond to the number of predicted target genes connected to each TF, and edge color intensity indicates interaction strength. (E) Literature-support summary for the top 10 TFs in the cattle lung and spleen GRNs. Each circle represents one TF among the top 10; circles are grouped by evidence level. Filled black circles indicate direct tissue-relevant support, filled gray circles indicate indirect support, and open circles indicate unsupported candidates under the current curation criteria. Detailed evidence is provided in Supplementary Table 4.

The matched-data species produced regulatory outputs that could be inspected in the portal without requiring each organism to have zebrafish-level coverage. In Mexican tetra, ELS sharing was strongest among liver samples from different populations, consistent with the shared organ identity of those samples (Supplementary Figure 13A) (64). In lancelet development, the largest number of unique ELSs appeared at 15 hpf, coinciding with somite formation (Supplementary Figure 13B) (65, 66). In Hydra regeneration, 485 ELSs were active across multiple regeneration stages and absent from the control condition, providing candidate regulatory elements associated with regeneration-state chromatin activity (Supplementary Figure 13C) (67). These analyses are framework outputs for matched datasets; ELS abundance was not interpreted as a direct cross-species biological comparison.

The cattle implementation provided an additional GRN example outside zebrafish. Motif enrichment in cattle liver ELSs recovered liver-associated factors including SP1, HNF4A, and PPARA (Figure 7B) (68–70). Other tissue-specific ELS sets also recovered tissue-relevant annotations: hypothalamus and cerebral cortex ELSs were enriched for neural TF motifs, adipose ELSs for adipocyte- and metabolism-related factors, and cattle cerebellum- or skeletal-muscle-associated genes for neural or muscle development terms (Supplementary Figures 13D, 14 and 15) (71–76). We also constructed cattle lung and spleen GRNs and curated literature support for the top ten TFs in each network (Figure 7C–E and Supplementary Table 4). In the lung GRN, three of the top ten TFs had direct lung-relevant support, including *AHR, ETS1*, and *YY1*; five had indirect support and two lacked sufficient support in our curation. Representative direct evidence includes endothelial AHR activity in lung barrier protection, Ets-1 involvement in developmental HIMF expression in mouse lung, and YY1 requirement for lung branching morphogenesis (77–79). In the spleen GRN, four of the top ten TFs had direct spleen- or immune-cell-relevant support, including *KLF3, STAT1, STAT3*, and *YY1*; the remaining six had indirect support. Representative direct evidence includes KLF3 programming of splenic marginal-zone B-cell fate, STAT1-dependent splenic macrophage trained immunity, STAT3-related spleen-cell responses, and YY1 requirement for germinal-center B-cell development (80–83). The cattle results provide a second GRN support summary outside zebrafish while keeping the interpretation at the level of candidate regulator prioritization.

The partial-data species illustrate the same framework under lower data completeness. Killifish was integrated as a new species during portal expansion and now has a Hugo species page, Dash gene information and bulk RNA modules, genome-browser tracks, SequenceServer databases, 1,348 bulk runs, and 31 single-cell sample groups comprising 386,393 cells. Because killifish uses local gene identifiers, its module depends on the species-adapter layer rather than changes to the shared API. Medaka uses a different partial-data configuration, including expanded bulk RNA coverage and dual identifier handling. These cases show that new species do not need to reach full miniENCODE core-assay completeness before the portal can provide useful access to available data.

This is the tiered output model used by miniODP. Complete or near-complete core assay datasets can be processed into ELS-centered outputs, with GRNs added for selected samples; species with partial data can still be integrated into the same portal and expanded as new assays become available.

### Comparison with existing resources

We compared miniODP with related species-focused omics resources by resource type, reusability, and data scale (Table 1). Zebrafish already has mature community resources, including ZFIN as a curated knowledgebase (6), DANIO-CODE as a functional-genomics data coordination center and regulatory-element atlas (8, 9), and recent single-cell or multimodal developmental atlases such as DanioCell, ZSCAPE, and ZebraHub (10–12). These resources are important references for zebrafish biology and for benchmarking regulatory annotations. They mainly serve curated knowledge, datasets, or atlas views for zebrafish and related analyses.

Recently, species-focused omics databases have been emerging.

For instance, iFish integrates genome, transcriptome, epigenome, and proteome resources across a large number of fish species (14). DOO provides a multi-omics atlas and analysis toolkit for deep-ocean organisms (15). MedakaBase offers a genome resource for multiple *Oryzias* species (16). In our comparison, these resources are suitable references, as they all integrate multiple types of data and offer web-based access. Therefore, we investigate whether each resource provides reusable deployment, new-species onboarding, and processing workflows, and whether these workflows can be reused to build a similar portal by other communities.

On these comparison dimensions, miniODP occupies a different position. Its contribution is not in having the largest number of species or datasets. Instead, this framework provides a deployable portal architecture, species-level configuration, gene-ID adapters, standardized data directories, cross-module links, and a staged onboarding process for adding new organisms. Seven-species implementation shows that the same architecture could support regulatory outputs when matched RNA-seq, ATAC-seq, and H3K27ac data are available, and also support partial-data species through core portal modules.

## Discussion

The main issue miniODP tries to solve is not another organism-specific website. For many understudied organisms, public omics data exist, but they are scattered across archives, source studies with diverse file formats, gene identifier systems, and genome assemblies. There is no practical route to turn the data into a maintained resource, so that researchers could browse, query, compare, and visualize them. miniODP is designed to solve this resource-building problem.

This perspective distinguishes miniODP from stand-alone databases or atlas projects. Existing resources like DANIO-CODE, DanioCell, ZSCAPE, ZebraHub, ZFIN, iFish, DOO, and MedakaBase serve curated knowledge, organism-specific atlases, or multi-species data access for defined communities (6, 8–12, 14– 16). miniODP neither tries to replace these resources nor competes with them by data volume or species number. Its main role is to provide a reusable path for building a resource: a reduced assay backbone, curated public and project data, species-specific identifier handling, linked gene and genome views, sequence search, and a staged route for adding new organisms.

miniENCODE core assays serve the same purpose. They are not a smaller version of ENCODE, which uses a broader set of assays to annotate regulatory function (2); they define a practical minimum for regulatory resource building: RNA-seq provides expression, ATAC-seq defines accessible chromatin, and H3K27ac profiling helps prioritize active enhancer-like and promoter-associated states. Zebrafish ELS-expression benchmark supports this design: with a data matrix of 3 assays x 17 samples, genes near H3K27ac-supported ELSs show higher expression levels than genes near ATAC-only distal peaks, even though the effect size is modest, and the analysis did not fully distinguish between ATAC signal strength and H3K27ac contribution. The main value is to show that the reduced assay set provides stable additional information for ELS definition.

We treat these outputs as a way to rank and inspect candidates, not as validation by themselves. ELSs, ELS-to-gene linkages, GRNs, reporter assays, and TF literature summaries give users several ways to inspect predictions. Tissue- and stage-specific ELSs showed expected TF-binding motif and GO enrichments. Selected heart-associated ELSs showed candidate-level reporter support, and GRN top TFs had different levels of direct, indirect, or insufficient literature support. These pieces of evidence can help users decide which predictions have stronger support, and which are merely hypotheses. If any single regulatory element or TF-target relationship needs to be validated, rigorous perturbation assays, stable reporter lines, and deeper sampling are still necessary.

The seven-species implementation showed that incomplete data availability is very common in non-model organisms, and can still serve as a starting point. Species without large numbers of matched core assays, such as medaka and killifish, can still support gene information, expression views, chromatin tracks, single-cell modules, BLAST search, and links to external annotations. When matched assays are available, additional regulatory analyses can be performed, and even two or three focused studies can form a species module.

This design also brings some practical limitations. When tissue coverage, assay depth, genome annotation, and public-data availability are different, ELS counts, SE counts, and GRN coverage cannot be directly used in cross-species biological comparisons. Non-standard local gene IDs can be handled by species adapters, but careful curation and manual checking remain important.

The central idea is that an omics resource for understudied organisms does not need to start as a complete atlas. It can begin by waking dormant data in the public archives and turning them into a usable infrastructure that grows with the data, the technology, and the community.

## Data Availability

ATAC-seq and CUT&Tag data are deposited in the NCBI Gene Expression Omnibus (GEO) database under accession GSE207155 (https://www.ncbi.nlm.nih.gov/geo/query/acc.cgi?acc=GSE207155; reviewer token obmtikycxzunfgz). The miniODP portal is accessible at: https://tulab.genetics.ac.cn/miniodp/. Source code, documentation, and Docker/Compose files are available at: https://github.com/QTuLab/miniodp. Docker images are available at: https://hub.docker.com/u/qtulab. Demonstration files for data analysis and visualization are available through ScienceDB at: https://doi.org/10.57760/sciencedb.41502.

## Supplementary Data

Supplementary Figures 1–15 and Supplementary Tables 1–5 accompany this preprint.

## Acknowledgements

We thank Dr. Jing-Wei Xiong (Peking University) for providing the TU strain of zebrafish; Dr. Tong Chen (State Key Laboratory for Quality Ensurance and Sustainable Use of Dao-di Herbs) and Mr. Minghong Zhang (Institute of Genetics and Developmental Biology) for assistance with website development.

AI-assisted tools were used for code development and language editing during manuscript preparation. All code, analyses, outputs, scientific interpretations, and final manuscript text were reviewed and verified by the authors, who take full responsibility for the work.

## Author contributions

Q.T. and Y.L. conceived and supervised the project. H.Y. performed most experiments and computational analyses. Z.W. contributed to miniODP implementation, data processing, and curation. Z.S. and H.S. contributed to the reporter assays. P.J. contributed to data processing and curation. Q.T., H.Y., and Y.L. wrote the manuscript with input from all authors.

